# Comparative analysis of beneficial effects of Vancomycin treatment on Th1- and Th2-biased mice and role of gut microbiota

**DOI:** 10.1101/516898

**Authors:** Pratikshya Ray, Uday Pandey, Palok Aich

## Abstract

**Aims:** Vancomycin, an antibiotic, is used to treat infection of multi-drug resistant strains of *Clostridium difficile* and *Staphylococcus*. Post-usage effects of vancomycin may lead to many unwanted effects including perturbation of gut microbiota. Perturbation of the gut microbiota, by Vancomycin, was used to understand the altered metabolic and innate immune profile of C57BL/6(Th1-biased) and BALB/c (Th2-biased) mice.

**Methods and Results:** Following treatment with vancomycin till day 4, we observed reduction in abundance of phyla Firmicutes and Bacteroides and increase in Proteobacteria in the gut for both strains of mice. Results further revealed a significant increase in the phylum Verrucomicrobia, from day 5 onwards following treatment with vancomycin led to decreased inflammation and increased rate of glucose tolerance in the host.

**Conclusions:** Continued dosage of vancomycin was more beneficial in C57BL/6 than BALB/c mice

**Significance and Impact of the study:** The current study established that initial doses of vancomycin increased pathogenic bacteria but the continued doses of vancomycin provided significant health-related benefits to the host by decreasing pathogenic load and by increasing beneficial microbes of Verrucomicrobia phylum (*A. muciniphila*) more in C57BL/6 (Th-1) than BALB/c (Th-2) mice.

## Introduction

More reports have started revealing that gut microbiota play an important role in maintaining health (Maldonado *et al*., 2012; Jandhyala *et al*., 2015; Andoh, 2016). Perturbation of gut microbiota can be used as an effective tool to understand its role in the host (Willing, Russell and Finlay, 2011). Abundance and diversity of gut microbiota change with different factors like age (Hill *et al*., 2017), diet (Singh *et al*., 2017), geography (Morton *et al*., 2015; Falony *et al*., 2016), stress (Bendtsen *et al*., 2012), pathogen (Khan, 2014), and antibiotics (Lange *et al*., 2016). Since treatment with antibiotics is still the most important and major avenue of dealing with many diseases, it is reported that the antibiotics could perturb the general taxonomy of the abundant and diverse gut microbes. Antibiotics are, therefore, being used as one of the most potent agents to study the role of gut microbiota (Jernberg *et al*., 2010). The altered microbial profile may lead to different diseases including metabolic syndromes like diabetes, obesity, and Inflammatory Bowel Disease (Bosi *et al*., 2006; Dunlop *et al*., 2006). Among various antibiotics, vancomycin can cause drastic changes in the human gut microbiota by increasing pathogens and by decreasing the commensal healthy microbes (Isaac *et al*., 2016). Vancomycin is majorly prescribed orally against the infection of two multi-drug resistant strains; i.e., *Clostridium difficile* and *Staphylococcus* (Bernard *et al*., 2003; Pepin, 2008; Tang *et al*., 2015). Reports also revealed that vancomycin could reduce gut microbial diversity and adversely affect metabolism and immunity of host (Vrieze *et al*., 2014; Isaac *et al*., 2016; Reijnders *et al*., 2016). The kinetics and dynamics of treatment with vancomycin on gut microbiota, however, are yet to be established. Especially, if treatment with vancomycin could affect the gut microbiome of Th1 and Th2-biased individuals differently to alter host metabolism and innate immunity. There were some earlier reports that elucidated the intrinsic differences in immune and immune-metabolic responses between C57BL/6 (Th1-biased) and BALB/c (Th2-biased) mice (Watanabe *et al*., 2004; Jovicic *et al*., 2015). The current laboratory reported earlier the differences in the abundance and composition of the gut microbiota between the two strains (Th1- and Th2-biased) of mice (Pradhan *et al*., 2018). Gut microbiota composition and diversity significantly influenced the immune system, gut barrier integrity and SCFA production of the host (Cani and Delzenne, 2009; Turner, 2009; Ulluwishewa *et al*., 2011; Wu and Wu, 2012). In this current study, we reported the role of vancomycin on the gut microbiota of both Th1- and Th2-biased mice. Time-dependent increase in specific gut microbes like Proteobacteria and Verrucomicrobia might be associated with the alteration of immune regulation, gut barrier maintenance, glucose metabolism and SCFA production of the host. The alteration pattern of gut microbes through vancomycin treatment and its effect on the host could be varied significantly between BALB/c and C57BL/6 mice. The results from the present study revealed that the earlier part of the vancomycin treatment caused i) an increase in abundance of specific pathogens, and ii) a decrease in native microbes. In the later stage of the treatment, some beneficial microbes started appearing and pathogens started declining, which provided health benefits to the host.

## Material and methods

### Animals Used in the study

All mice of the same strain used in the present study were co-housed in polysulfone cage, and corncob was used as bedding material. Food and water were provided *ad libitum*. Animals were co-housed in a pathogen-free environment with a 12h light-dark cycle at temperature 24 ± 3° with the humidity around 55%. The guidelines for animal usage were as per CPCSEA (Committee for the Purpose of Control and Supervision of Experiments on Animals, Govt. of India). All protocols were approved by the Institute Animal Ethics Committee constituted by CPCSEA (Reg. No.- 1643/GO/a/12/CPCSEA). 6-8 weeks old male C57BL/6 (Th1) and BALB/c (Th2) mice were used in the present study. Unless otherwise stated, we used at least 3 mice per treatment condition per time point.

### Antibiotic treatment

Both Th1-(C57BL/6) and Th2- (BALB/c) biased mice were treated (n=3 per group) with vancomycin (Cat#11465492) twice daily for 6 consecutive days. Vancomycin was administered by oral gavaging at 50 mg per kg of body weight. The dosage was selected as per previous reports and FDA guidelines(Erikstrup *et al*., 2015; Patel, Preuss and Bernice, 2019)

### Mice treatment and sample collection

Mice were separated into two different groups: Control (untreated) and Treatment (groups that were treated with vancomycin). Each day of the experiment for 6 days, time-matched control and treated mice were euthanized by cervical dislocation. Colon tissue, serum and cecal materials were isolated from each mouse following the protocols described elsewhere. Tissue samples, not used immediately, were stored (−80°C) either by snap freezing for protein analysis or in RNAlater for RNA analysis until further used (Pradhan *et al*., 2018; Ghosh *et al*., 2019)

### Cecal Sample plating

Cecal sample was collected from both strains of mice on day 4 following treatment with vancomycin. 50 mg of each sample was homogenized in 1ml of deionized MilliQ water and plated at a dilution of 10^4^ fold on Salmonella-Shigella specific media and EMB (Eosin methylene blue agar plate) agar plate (Dekker and Frank, 2015).

### RNA extraction

RNA was extracted from the colon tissue by using RNeasy mini kit (Cat# 74104, Qiagen, Germany) following the manufacturer’s protocol. 20-23 mg of tissue was processed using liquid nitrogen followed by homogenization in 700 μl of RLT buffer. An equal volume of 70% ethanol was added and mixed well. The solution was centrifuged at 8000g for 5 minutes at room temperature. The clear solution containing lysate was passed through RNeasy mini column (Qiagen, Germany), which leads to the binding of RNA to the column. The column was washed using 700 μl RW1 buffer and next with 500 μl of RPE buffer. RNA was eluted using 30 μl of nuclease-free water. RNA was quantified using NanoDrop 2000 (ThermoFisher Scientific, Columbus, OH, USA).

### cDNA preparation

cDNA was synthesized by using Affinity Script One-Step RT-PCR Kit (Cat# 600559, Agilent, Santa Clara, CA, USA). RNA was mixed with random nonamer primers, Taq polymerase, and NT buffer. The mixture was kept at 45 °C for 30 min for the synthesis of cDNA and temperature increased to 92 °C for deactivating the enzyme.

### Real-time PCR (qRT-PCR)

Real-time PCR was performed in 96-well plate, using 25 ng cDNA as template, 1 μM of each of forward (_F) and reverse (_R) primers for genes mentioned in Table 2, SYBR green master mix (Cat#A6002, Promega, Madison, WI, USA), and nuclease-free water. qRT-PCR was performed in Quantstudio 7 (Thermo Fisher Scientific, Columbus, OH, USA). All values were normalized with cycle threshold (Ct) value of GAPDH (internal control) and fold change of the desired gene was calculated with respect to the control C_t_-value as mentioned elsewhere (Pradhan *et al*., 2016, 2018).

**Table 1.**
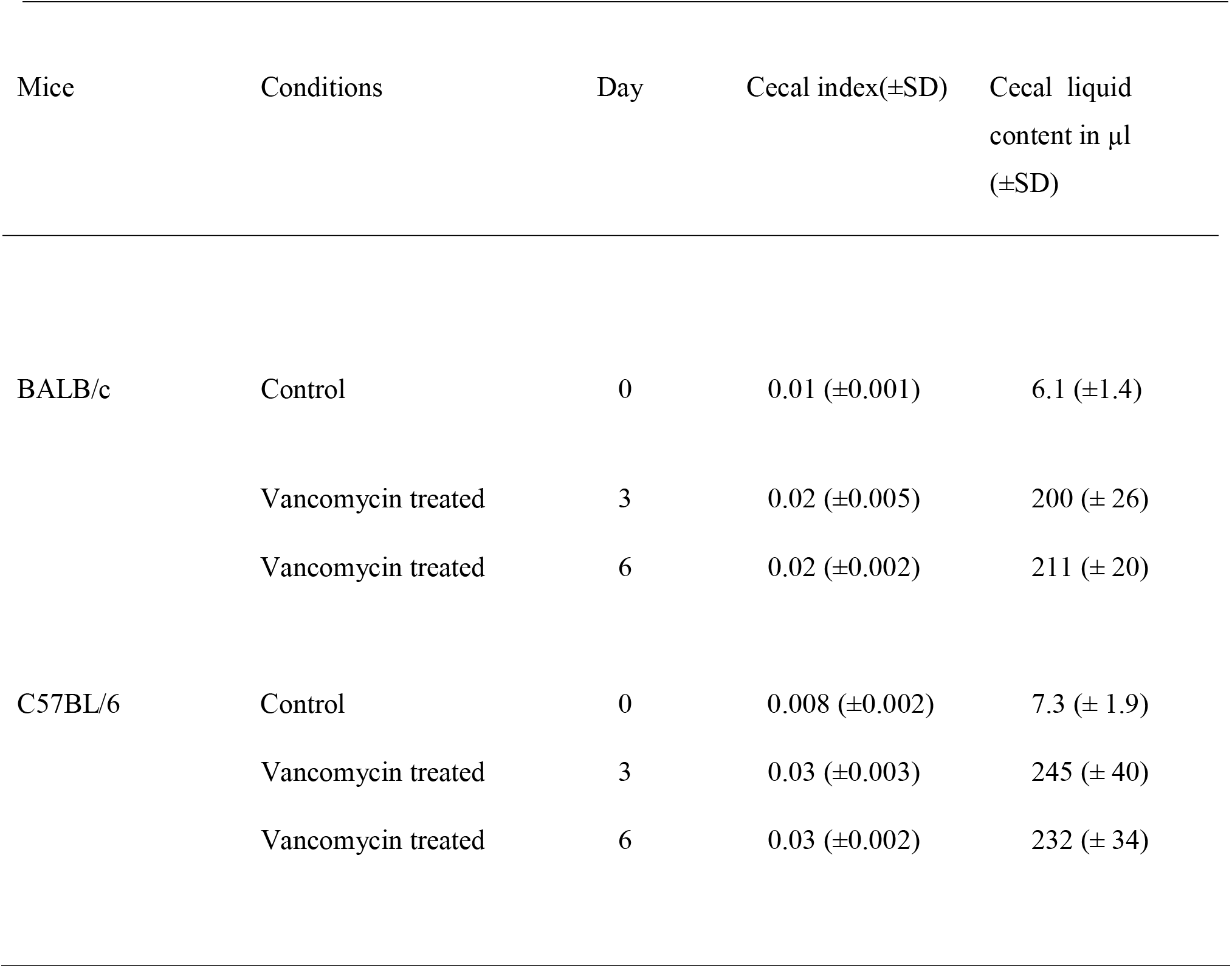
Measurement of Cecal index and cecal liquid content at different time points of BALB/c and C57BL/6 mice.

**Table 2.**
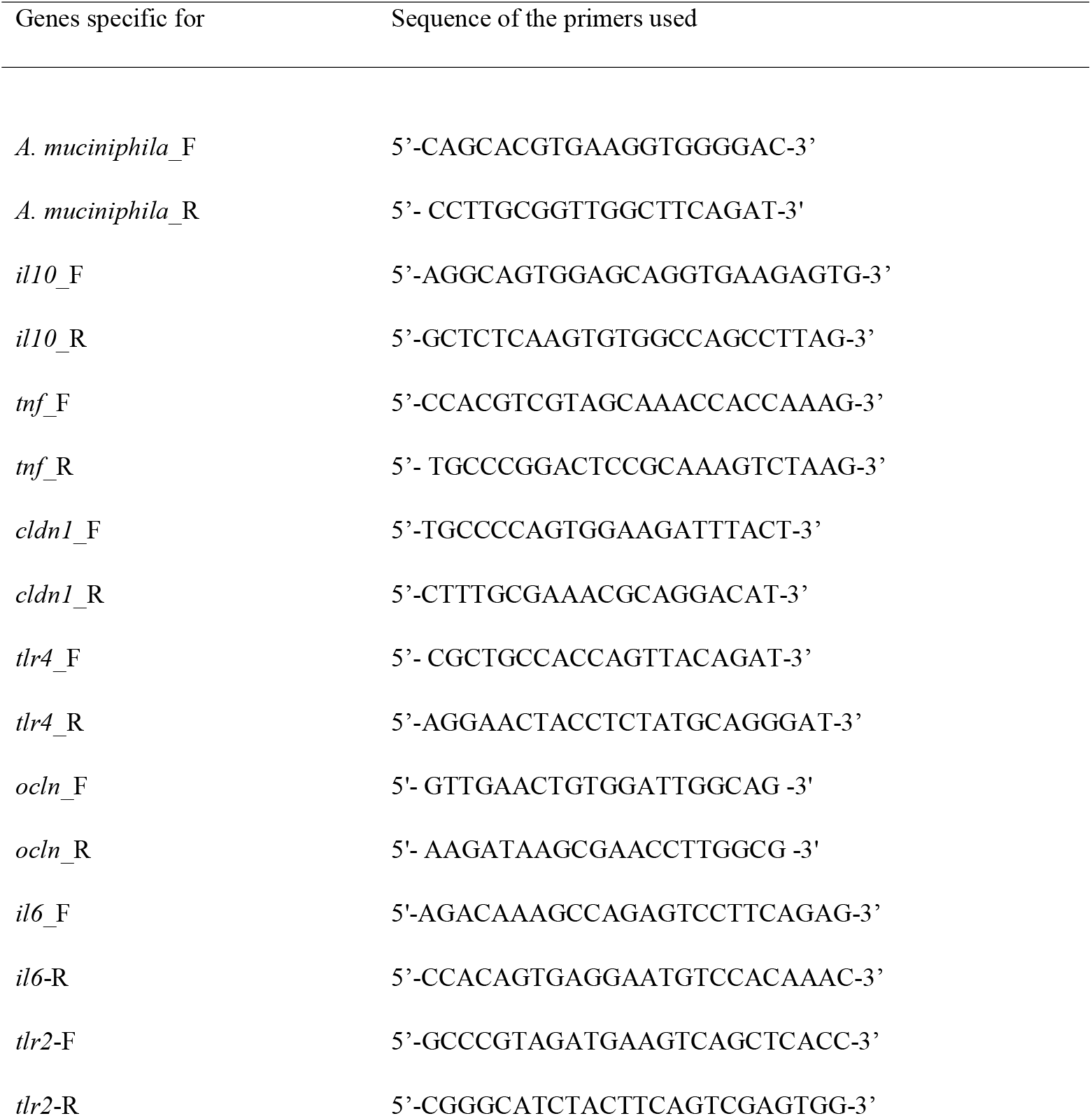
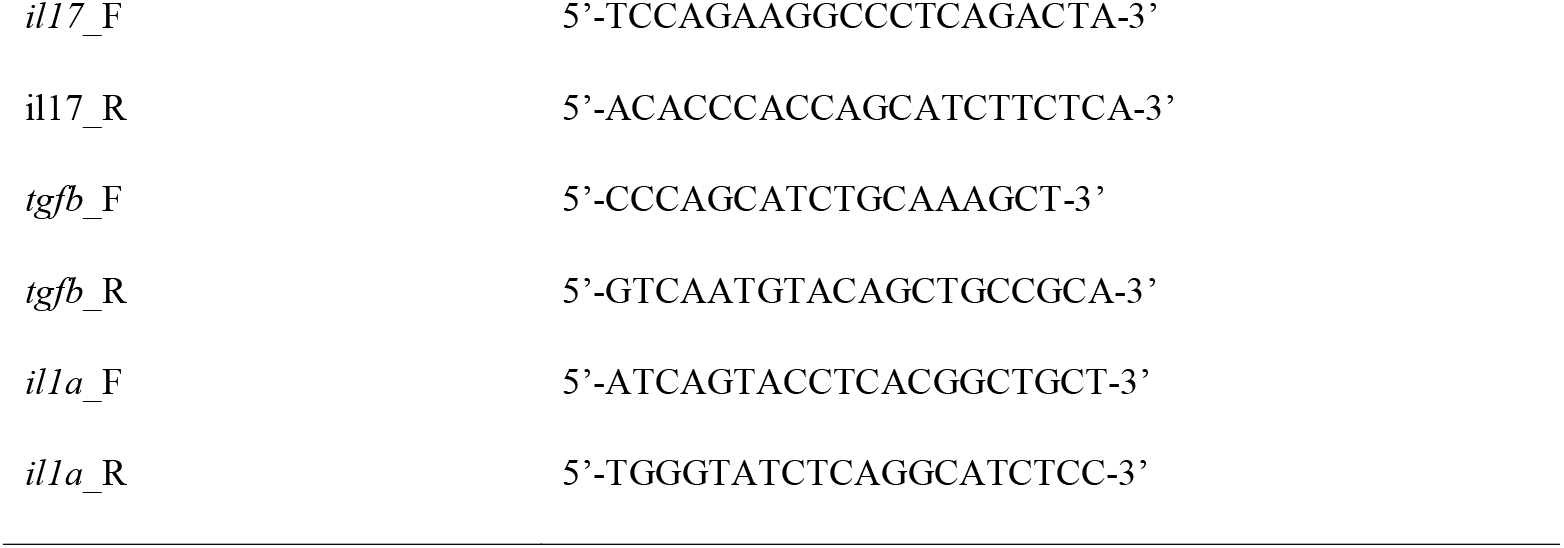
Sequences of forward (_F) and reverse (_R) primers for PCR studies to confirm presence and expression level of various genes used in this study.

### Cytokine Analysis at the protein level

Colon tissues were collected from mice on day 0 (untreated control) and on days 3 and 6 following treatment with vancomycin. After washing the colon tissues thoroughly, lysis buffer (Tris-hydrochloric acid, sodium chloride, and Triton X-100 in distilled water) containing 1X protease inhibitor cocktail (PIC) (Cat#ML051, Himedia, India) was used to churn the tissue [40]. The supernatant was collected following centrifuging the churned mixture at 20,000g for 20 minutes. ELISA (BD Biosciences, San Diego, CA, USA) was performed using the manufacturer’s protocol for TNFα (Cat#560478) and IL10 (Cat#555252) expression [40]. Protein concentration was normalized through Bradford assay (BD Biosciences, San Diego, CA, USA). The absorbance was taken using Multiskan Go (Thermo Fisher Scientific, Columbus, OH, USA).

### Serum collection

Mice were anaesthetized and whole blood was collected by cardiac puncture. Blood was kept on ice for 30 mins and centrifuged at 1700g for 15 min at 4°C, and serum was collected for further analysis. If required, serum was stored at −80°C until further use.

### Genomic DNA extraction

Cecal sample was collected from mice of both strains and gDNA was extracted using the phenol-chloroform method. 150-200 mg of cecal sample was used to homogenize using 1ml of 1X PBS and centrifuged at 6700g for 10 minutes. The precipitate was lysed by homogenizing it in 1ml of lysis buffer (containing Tris-HCl 0.1M, EDTA 20 mM, NaCl 100 mM, 4% SDS (at pH 8) and thereafter heating it at 80 °C for 45min. Lipid and protein were removed from the supernatant using an equal volume of phenol-chloroform, this process was repeated until the aqueous phase became colourless. DNA was precipitated overnight at −20 °C with 3 volumes of absolute chilled ethanol. Finally, it was washed with 500 μl of 70% chilled ethanol and briefly air-dried. The gDNA was dissolved in nuclease-free water and quantified using NanoDrop 2000.

### 16S-rRNA sequencing (V3-V4 Metagenomics)

Using cecal DNA samples, V3-V4 regions of the 16S rRNA gene were amplified. For this amplification, V3F (Forward primer): 5’-CCTACGGGNBGCASCAG-3’ and V4R (Reverse primer): 5’-GACTACNVGGGTATCTAATCC-3’ primer pair was used. In Illumina Miseq platform, amplicons are sequenced using paired-end (250bpX2) with a sequencing depth of 500823.1 ± 117098 reads (mean ± SD). Base composition, quality and GC content of FASTQ sequence were checked. More than 90% of the sequences had Phred quality scores above 30 andGC content nearly 40-60%. Conserved regions from the paired-end reads were removed. Using the FLASH program, a consensus V3-V4 region sequence was constructed by removing unwanted sequences (Kim *et al*., 2012; Mysara *et al*., 2017). Pre-processed reads from all the samples were pooled and clustered into Operational Taxonomic Units (OTUs) by using de novo clustering method based on their sequence similarity using UCLUST program. QIIME was used for the OTU generation and taxonomic mapping (Caporaso *et al*., 2010; Purcell *et al*., 2017). A representative sequence was identified for each OTU and aligned against the Greengenes core set of sequences using the PyNAST program(DeSantis *et al*., 2006b, 2006a; Frank *et al*., 2007; Kastenberger *et al*., 2012). Alignment of these representative sequences against reference chimeric data sets was done and the RDP classifier against the SILVA database was used for taxonomic classification to get rid of hybrid sequences.

### Cecal Microbiota Transplantation (CMT)

Cecal sample was collected from sixth-day vancomycin treated mice and diluted with PBS (1gm per10ml) to make stock. 400 μl of the stock of cecal material was orally gavaged to each of third-day vancomycin treated mice.

### Oral Glucose tolerance test (OGTT)

OGTT was assayed on days 0, 3 and 6 following the treatment of both types of mice with vancomycin. Following 6h of starvation of mice from each treatment group, fasting blood glucose level (considered as control glucose level at 0 minutes) was measured by a glucometer by tail vein bleeding. Mice, fasted for 6h, were orally gavaged with glucose at a dose of 1 mg g^-1^ bodyweight of the mouse. Blood glucose levels were measured at intervals of 15-, 30-, 60- and 90-minutes post-glucose gavaging by using a blood glucose monitoring system (ACCU-CHEK Active, Roche Diabetes Care GmbH, Mannheim, Germany).

The same procedure was adopted for oral glucose tolerance test was done for third day CMT recipient mice following 24h of CMT procedure.

### Sample preparation and NMR data acquisition for metabolomics study

Serum was isolated from the blood of vancomycin treated and control mice as described before. Proteins in the serum were removed by passing it through a pre-rinsed (7 times washed) Amicon Ultra-2ml 3000 MWCO (Merck Millipore, USA) column. Centrifugation was done at 4°C at 12,000g. Total of 700 μL solution (containing serum sample, D_2_O, pH maintenance buffer and DSS) was taken in 5 mm Shigemi tubes. NMR for all samples were performed at 298K on a Bruker 9.4 T (400 MHz) AVANCE-III Nanobay liquid-state NMR spectrometer equipped with 5 mm broadband (BBO) probe. The pre-saturation technique was used with a moderate relaxation delay of 5 seconds to ensure complete water saturation. Offset optimization was performed using a real-time ‘gs’ mode for each sample. Topspin 2.1 was used to record and process the acquired spectra.

### Metabolomic Analysis of NMR data

ChenomX (Canada) was used for the analysis of NMR data. The Bayesian approach is used to derive metabolite concentration in serum. The phase and baseline of the raw spectrum were corrected and concentrations of metabolites were obtained through a profiler using Metaboanalyst (Hapfelmeier *et al*., 2005; DeSantis *et al*., 2006a; Frank *et al*., 2007; Xia and Wishart, 2011; Xia *et al*., 2015; Zhou and Zhi, 2016). To normalize the data across the study, the samples were log-transformed and compared with the control sample. Relative fold-change values in metabolite expression analysis were performed for each treated samples with respect to the untreated time-matched control. More than 2-Fold change values (above or below reference value) with p□ ≤ 0.05 were considered for further analysis.

### Calculation of Cecal index

The body weight of each individual mouse was measured and recorded. The whole cecal content was collected in a microfuge tube and weighed for each individual mouse. The cecal index was measured by taking the ratio of cecal content to the body weight of each mouse and used by normalizing the data with respect to the average body weight of mice used(Shi *et al*., 2018).

### Gut permeability test by FITC dextran

Gut permeability was determined by measuring the concentration of non-digestible dextran conjugated with fluorescein isothiocyanate in the serum. After oral administration, FITC dextran transits through the GI tract and crosses the intestinal epithelium. Mice used for this experiment were water-starved overnight. Next day morning, FITC-dextran (Cat#F7250, Sigma-Aldrich, Missouri, US) at a concentration of 100mg ml^-1^ was dissolved in PBS and oral gavaged. After 4h, mice were anaesthetized by isoflurane inhalation and the blood was collected by cardiac puncture. Serum was collected from blood and concentration of FITC in serum was measured by spectrofluorometer (Varioskan, ThermofisherScientific) with an excitation wavelength of 485 nm (20 nm bandwidth) and emission of 528 nm (20 nm bandwidth)(Woting and Blaut, 2018).

### Endotoxin detection assay from serum

Limulus Amebocyte Lysate (LAL) test was used for the detection of lipopolysaccharides located in the outer cell membrane of gram-negative bacteria. For this test, mice were sacrificed on days 0, 3 and 6 following treatment of mice with vancomycin and blood was collected by cardiac puncture in an endotoxin-free vial. Toxin sensor chromogenic LAL endotoxin assay kit from GeneScript (Cat#L00350, Piscataway, NJ, USA) was used for detecting endotoxin level in the serum of mice using the manufacturer’s protocol (Holzheimer, 2014).

### Acetate detection assay in serum

Acetate level was measured in the serum samples of untreated (control day 0) and day 6 following treatment of BALB/c and C57BL/6 mice with vancomycin by using acetate colorimetric assay kit (EOAC-100, San Francisco, USA). Both control and treated mice were anaesthetized and blood was collected through cardiac puncture. Blood was kept on ice for 30 mins followed by centrifugation at 1700g for 15 min at 4 °C. The supernatant was collected and 10 μl of serum from each sample was used to detect acetate level using substrate-enzyme coupled colorimetric reaction assayed by absorbance at 570 nm.

### Hormonal assay

Leptin (Cat# ELM-Leptin), and Insulin (Cat# ELM-Insulin) hormone levels were assayed in serum samples. PYY (Cat# EIAM-PYY) was assayed in the colon tissue samples by using Raybiotech mouse hormonal assay kit (Norcross, Georgia, USA).

### Statistical Analysis

All the graphs were plotted using GraphPad Prism version 7.0. Statistical package in Prism was used for statistical analysis for the data to perform a ‘t’-test (to compare any 2 data sets) or ANOVA (to compare more than two datasets) as described in the text.

## Results

### Vancomycin treatment alters the abundance and diversity of gut microbiota

Effective gut microbial perturbation by vancomycin treatment was previously reported (Isaac *et al*., 2016)but the role of the immune profile of the host was not addressed. The mammalian hosts could be broadly discriminated based on the immune profile in terms of pro-inflammatory (Th1) and tolerogenic (Th2) responses. We, therefore, compared the differential effects, if any, of vancomycin onTh1- and Th2-immune biased mice strains (C57BL/6 and BALB/c). Changes in the gut (specifically small and large intestine) associated physical and morphological parameters are usually the first important signs to look for following antibiotic treatment (Jernberg *et al*., 2010). We observed that the vancomycin treatment significantly increased the cecal index (an important parameter determined by cecum weight/body weight) in both strains of the mice. The increase in the cecal index also suggested the alteration of bacterial abundance in the cecum (Patel *et al*., 2012; Shi *et al*., 2018) (Table. 1). We used 16S rRNA (metagenomic) based sequencing protocol to understand the kinetics of altered microbiota profile in the cecum following treatment with vancomycin. Metagenomic analysis of the cecal content revealed that the microbial composition was significantly altered in both BALB/c and C57BL/6 mice following treatment with vancomycin (Figure 1). The results of untreated mice, shown in Figure 1A and Figure 1C, mainly revealed that in both BALB/c and C57BL/6 the gut microbiota overtly belong to the phyla, Firmicutes and Bacteroidetes. The abundance of the phyla, Firmicutes and Bacteroidetes, reduced while the abundance of Proteobacteria phylum increased significantly by the second day following treatment with vancomycin. The Proteobacteria level reached maximum by day five (93% of total abundance) in BALB/c mice and day four (81% of total abundance) in C57BL/6 following treatment with vancomycin (Figure 1B and Figure 1D). On the contrary, Firmicutes level decreased from 70-80% (untreated control group) to below 10% (fourth day following treatment with vancomycin) and Bacteroidetes level from 25-30 % (untreated group) to 1% (fourth day following treatment with vancomycin) in both BALB/c and C57BL/6 mice (Figure 1B and Figure 1D). After day 4 of treatment with vancomycin, BALB/c and C57BL/6 mice showed a significantly different gut microbiota profile with the appearance of phylum, Verrucomicrobia. A sudden increase in Verrucomicrobia phylum, from day five onwards, in C57BL/6 and from day six onwards, in BALB/c mice following vancomycin treatment replaced the previously predominant Proteobacteria phylum. However, the abundance of Verrucomicrobia phylum was found to be significantly higher, in C57BL/6 mice (72% of total abundance), than in BALB/c mice (30% of total abundance) on the day six following treatment with vancomycin. This result was significant to understand the differential response exhibited in two different strains of mice (C57BL/6 and BALB/c) used in this study following treatment with vancomycin.

**Figure 1.**
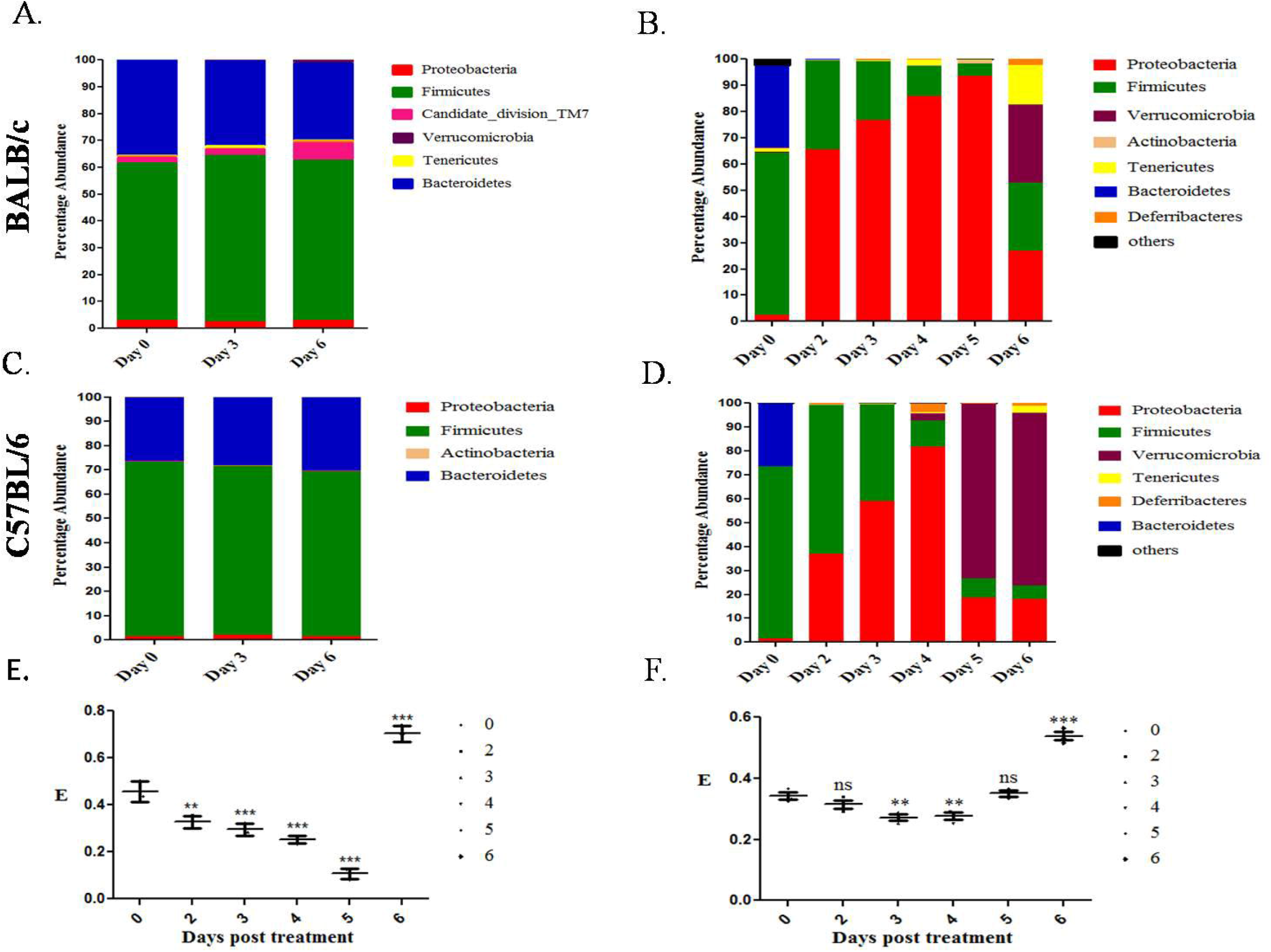
Phylum level changes in the gut microbiota. Time dependent changes in the phyla of gut microbiota, in the control A. BALB/c and C. C57BL/6 and in vancomycin treated B. BALB/c and D. C57BL/6 mice are shown. In the figure, only major phyla are shown to avoid clutter. Data shown are average of 3 biological replicates. To avoid clutter, standard deviation (SD) calculated using 2-way ANOVA is not shown. However, SD was less than 10% on average. Kinetics of changes in phylum level Equitability index (E) (diversity) of the gut microbiota following treatment with vancomycin in E. BALB/c and F. C57/BL6 mice. Statistical significance of diversity in panels E. And F. was calculated by two-way ANOVA. (‘***’ corresponds to P ≤ 0.001, ‘**’ corresponds to P ≤ 0.01, ‘*’ corresponds to P ≤ 0.05 level of significance). Error bars are one standard deviation of the mean value and determined from the average values of biological replicates.

In addition, we further determined the Shannon diversity of the cecal microbiota. Shannon equitability index at the phylum level showed a decrease in diversity up to the fifth day in BALB/c (Figure 1E) and up to the fourth day in C57BL/6 mice (Figure 1F) following treatment with vancomycin. Microbial diversity was increased on day 6 for BALB/c and day 5 for C57BL/6 following treatment with vancomycin.

Since each phylum contains various genera, we were interested to find out the changes in the abundance and diversity at the genus level following vancomycin treatment. At the genus level, the gut microbiota of untreated time-matched control mice majorly composed of Blautia, Intestinimonas genera of Firmicutes phylum and Bacteroides, Alistipes genera of Bacteroidetes phylum in both strains (Figure 2A and Figure 2C). Whereas, on day four, the gut microbiota of vancomycin treated mice showed mostly Escherichia-Shigella and Desulfovibrio genus from Proteobacteria phylum in both strains of mice(Figure 2B and Figure 2D). These results were further validated by plating day four cecal homogenate in specific media-EMB agar (*E.coli*) and Salmonella-Shigella agar plate (*Shigella sp.)*. Plating data of cecal samples from day four following vancomycin treatment showed overgrown colonies compared to untreated mice on the specific media (Figures 2E and 2F). On the day six following vancomycin treatment, the genus level data showed a predominance of *Akkermansia* in both strains of mice. However, *Akkermansia muciniphila* level was higher in C57BL/6 mice than BALB/c mice. We performed 16S based qPCR by using *Akkermansia muciniphila* species-specific primers to confirm metagenomic data (Table 4). The qPCR results revealed that there were nearly 21 fold and 24833 fold increase in *A. muciniphila* abundance in vancomycin treated BALB/c and C57BL/6 mice respectively on day six compared to their time-matched untreated control mice. While on day three following vancomycin treatment, due to very low abundance of *A. muciniphila*, the value of threshold cycle (Ct), from qPCR data, could not be determined for either of BALB/c or C57BL/6 mice. The change in the abundance of *A. muciniphila* was found to be comparable in both metagenomic and qPCR analysis.

**Figure 2.**
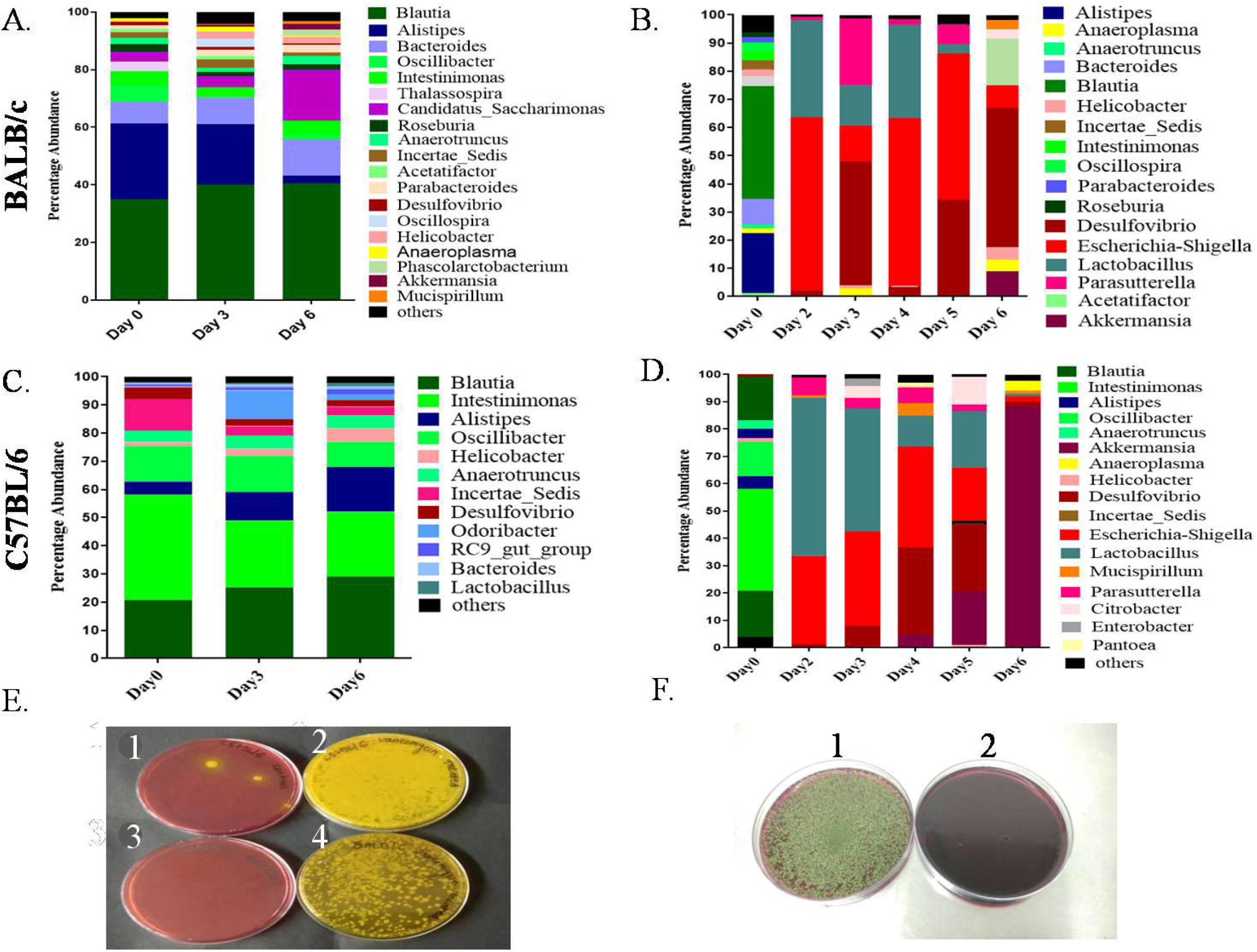
Metataxonomic studies of genus level variation of gut microbiota in vancomycin treated and its respective control group. Kinetics of changes in genera of gut microbiota, in the control A. BALB/c, and C. C57BL/6 and in vancomycin treated B. BALB/c, and D. C57BL/6 mice are shown. Data shown are average of 3 biological replicates and standard deviation was less than 10% on average. Percentage abundance of different genera for various treatment conditions are shown on the ‘Y’-axis and the days elapsed post treatment or for time matched control are shown on the ‘X’-axis. Colony of culturable Proteobacteria by plating of cecal samples from both strains of mice on selective and differential media. Evidence of E. Shigella colonies growth on day 4 on Salmonella-Shigella specific media agar plate 1. control C57BL/6, 2. vancomycin treated C57BL/6, 3. control BALB/c and 4. vancomycin treated BALB/c and F. Growth of *E. coli*. colonies on day 4 on EMB (Eosin methylene blue agar plate), 1. vancomycin treated C57BL/6 and 2. control C57BL/6.

**Table 3.**
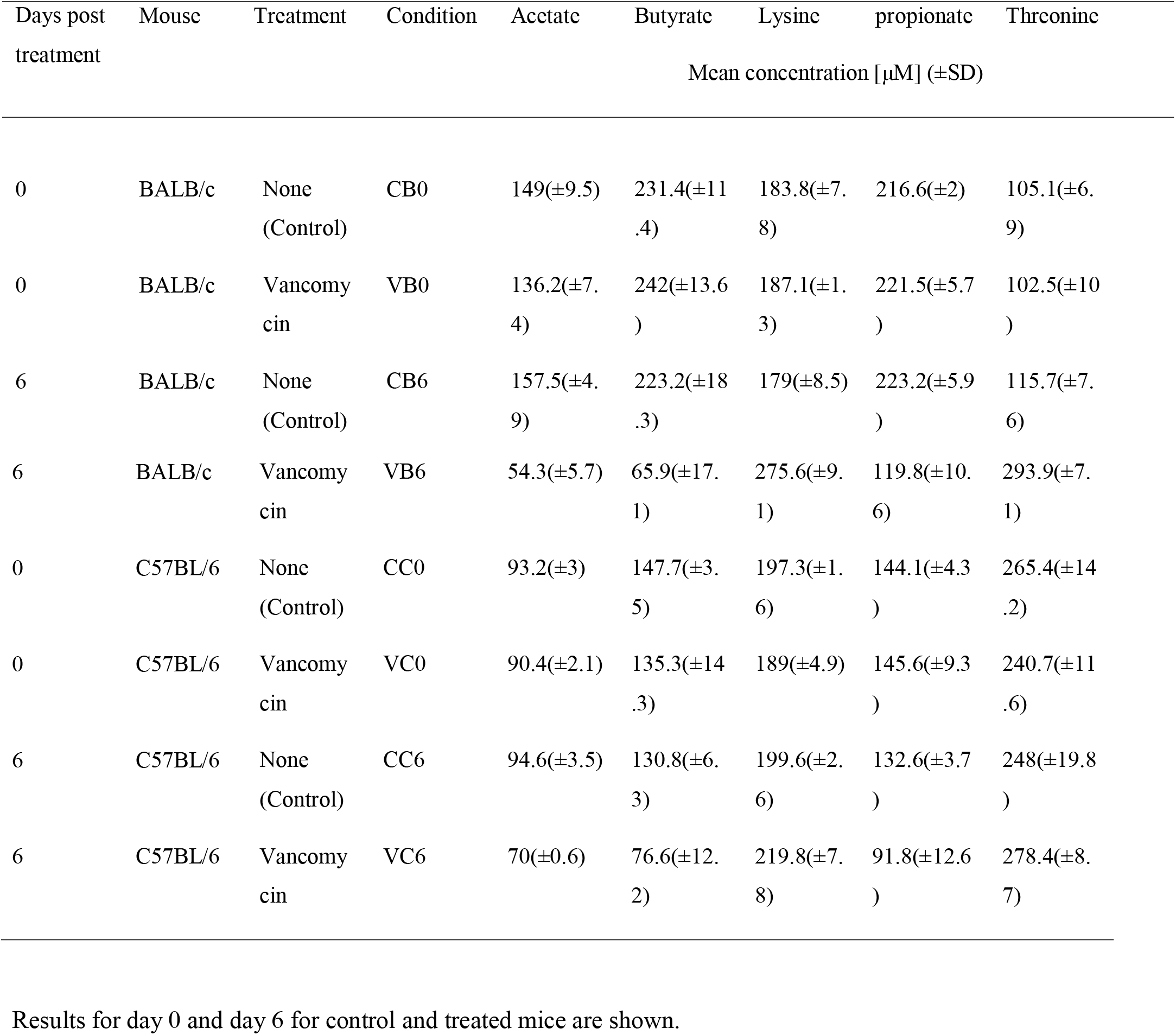
Abundance of various SCFAs and associated metabolites in untreated (control) and vancomycin treated BALB/c and C57BL/6.

**Table 4.**
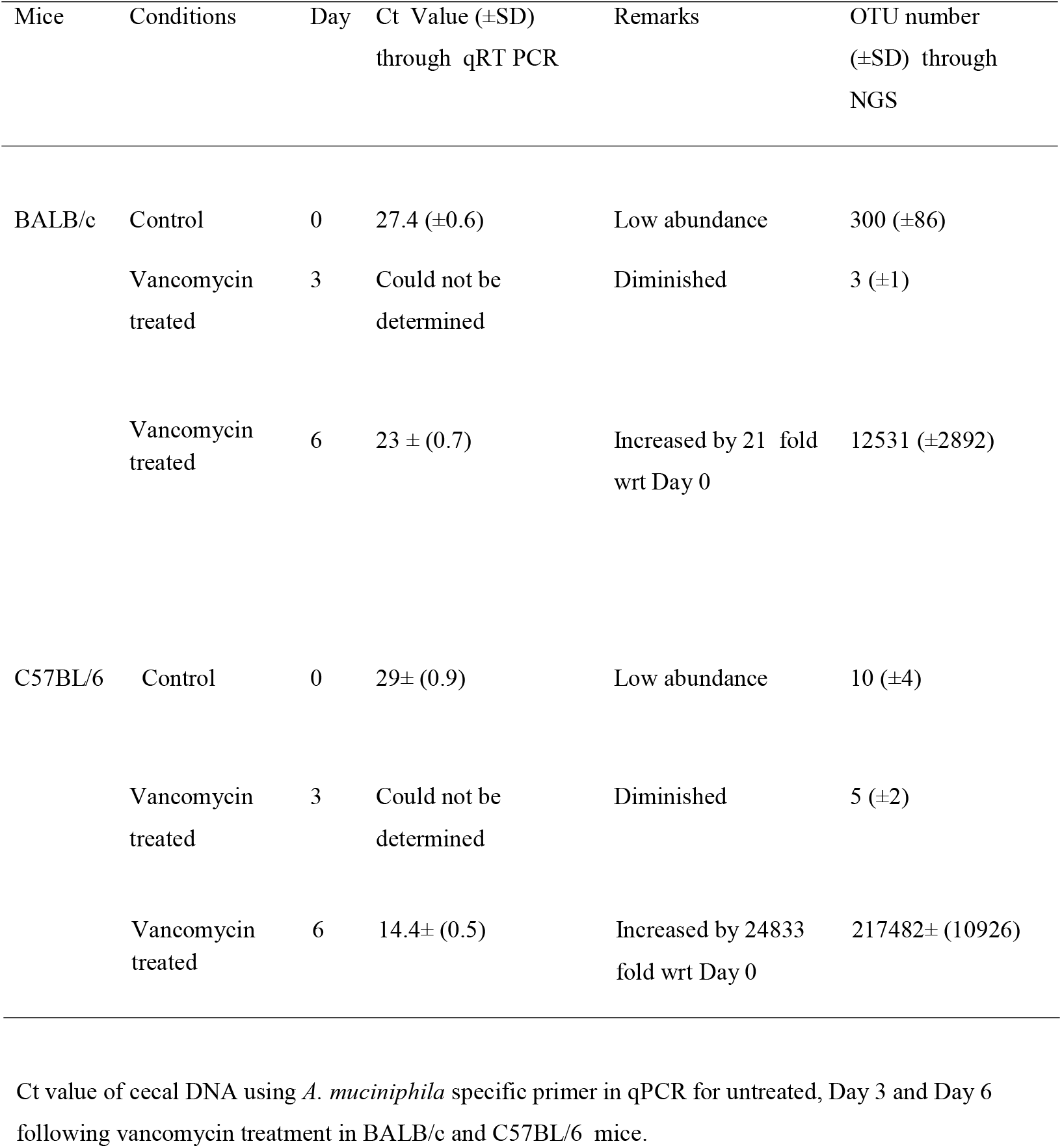
16S qPCR detection of *A. muciniphila* bacteria abundance in cecal sample of treated and control mice.

Above results indicated that both the mice strains showed an initial increase in Proteobacteria abundance following treatment with vancomycin in a time-dependent manner followed by an increase in abundance of Verrucomicrobia Phylum by day six. Also, the differential abundance of *A. Muciniphila* on day six showed a difference in the response of vancomycin perturbation of gut microbiota between two strains of mice.

### The inflammatory response in the colon changes during vancomycin mediated microbiota perturbation in a time-dependent manner

Antibiotic induced gut microbiota perturbation is associated with different inflammatory disorders (Willing *et al*., 2011; Miyoshi *et al*., 2017). We checked the effect of vancomycin mediated microbial perturbation on the expression of various pro-(tnfα, il6, il1a, il17) and anti-inflammatory (tgfβ and il10) genes in both mice strains. The mRNA level expression data from colonic tissue revealed the time-dependent increase of pro-inflammatory cytokines till day four of treatment in both mice strains (Figures 3A and 3B). A decrease in the expression of the pro-inflammatory cytokines was associated with the decrease of Proteobacteria abundance after day four of vancomycin treatment (Figures 3C and 3D). Next, we also observed a marked increase of tlr4 expression, upstream regulator of the inflammatory response (Wahid *et al*., 2015) on the third day and decreased by the sixth day of treatment in both mice strains (Figures 3E and 3F). However, we found a significant increase in the expression of tlr2 on day five and day six following vancomycin treatment in C57BL/6 mice compared to BALB/c mice. The increase of tlr2 gene expression was correlated with the higher abundance of *A. muciniphila* during the day five and day six following vancomycin treatment in C57BL/6 mice. Validation of qRT-PCR results was done, at the protein level expression by ELISA (Figures 4A and 4C). ELISA results revealed that on the third day following vancomycin treatment, TNFα level was significantly more in both BALB/c and C57BL/6 mice with respect to the third day time-matched untreated groups of mice. Similarly, IL10 cytokine level was more in BALB/c and C57BL/6 on day six following treatment with vancomycin compared to the day sixth time-matched untreated mice (Figures 4A and 4C). Both qRT PCR and ELISA data showed nearly similar results for the expression of pro- and anti-inflammatory cytokines (Figures 4B and 4D). In summary, our data suggest that the pro-inflammatory response in colonic tissue is linked with increased Proteobacteria abundance during vancomycin mediated microbial disruption. The emergence of Verrucomicrobia phyla from the fifth day onwards may lead to a transition from pro-inflammatory to anti-inflammatory response irrespective of initial immune bias of the mice. However, on the sixth day following vancomycin treatment, the decrease of pro-inflammatory cytokine and an increase of anti-inflammatory cytokine expression was more significant in C57BL/6 mice compared to BALB/c mice. This result can be correlated with the difference in the abundance of Verrucomicrobia phylum between BALB/c and C57BL/6 mice on the sixth day following vancomycin treatment.

**Figure 3.**
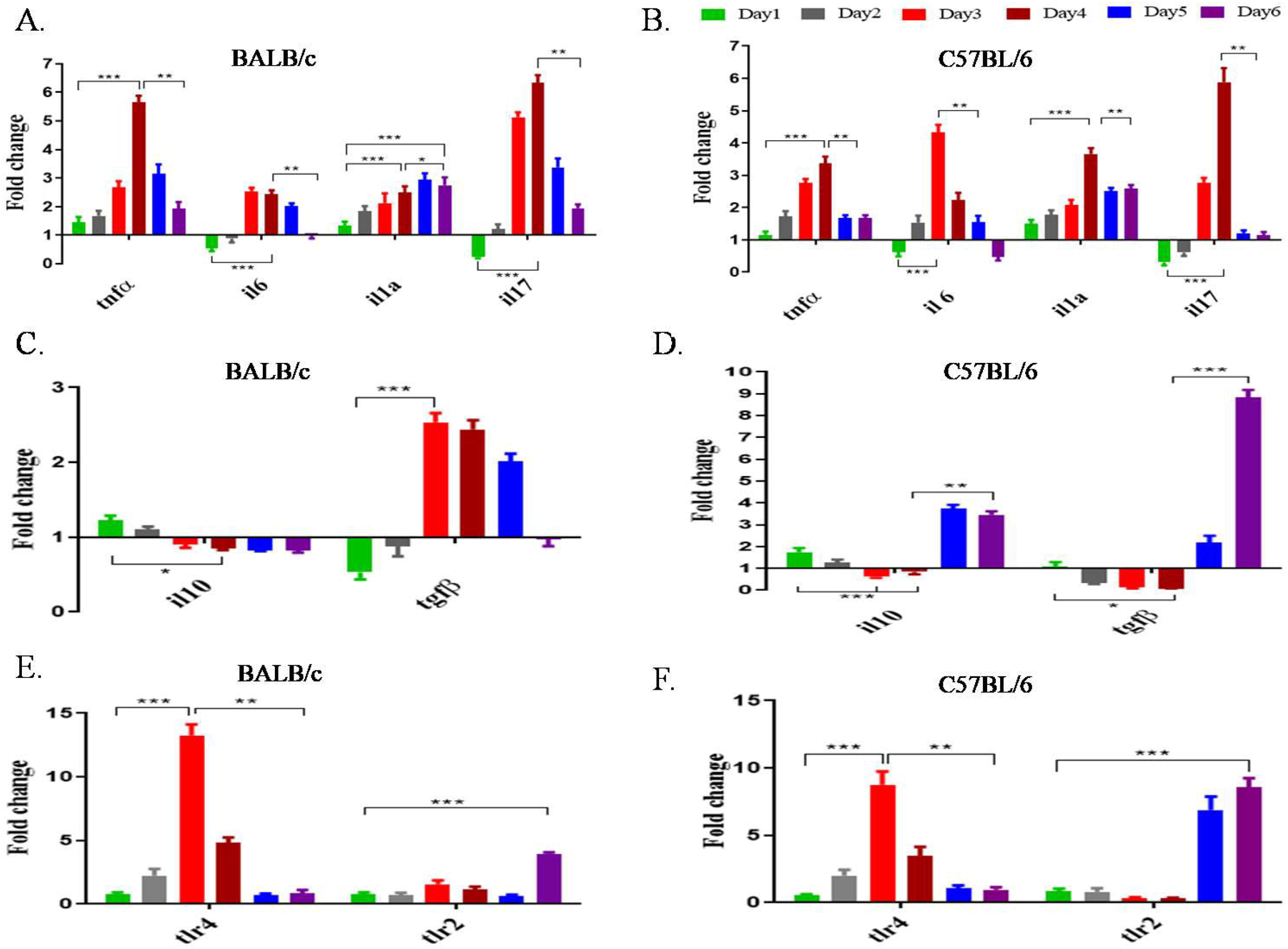
Transcriptional gene expression profile. Kinetics of transcriptional (by qRT-PCR) expression of genes categorized as, pro-inflammatory in A. BALB/c, and B. C57BL/6, anti-inflammatory in C. BALB/c, and D. C57BL/6 as Toll like receptors tlr4 and tlr2 in E. BALB/c, and F. C57BL/6 mice. Statistical significance was calculated by two-way ANOVA. (‘***’ corresponds to P ≤ 0.001, ‘**’ corresponds to P ≤ 0.01, ‘*’ corresponds to P ≤ 0.05 level of significance). Error bars are one standard deviation of the mean value and determined from the average values of three biological replicates.

**Figure 4.**
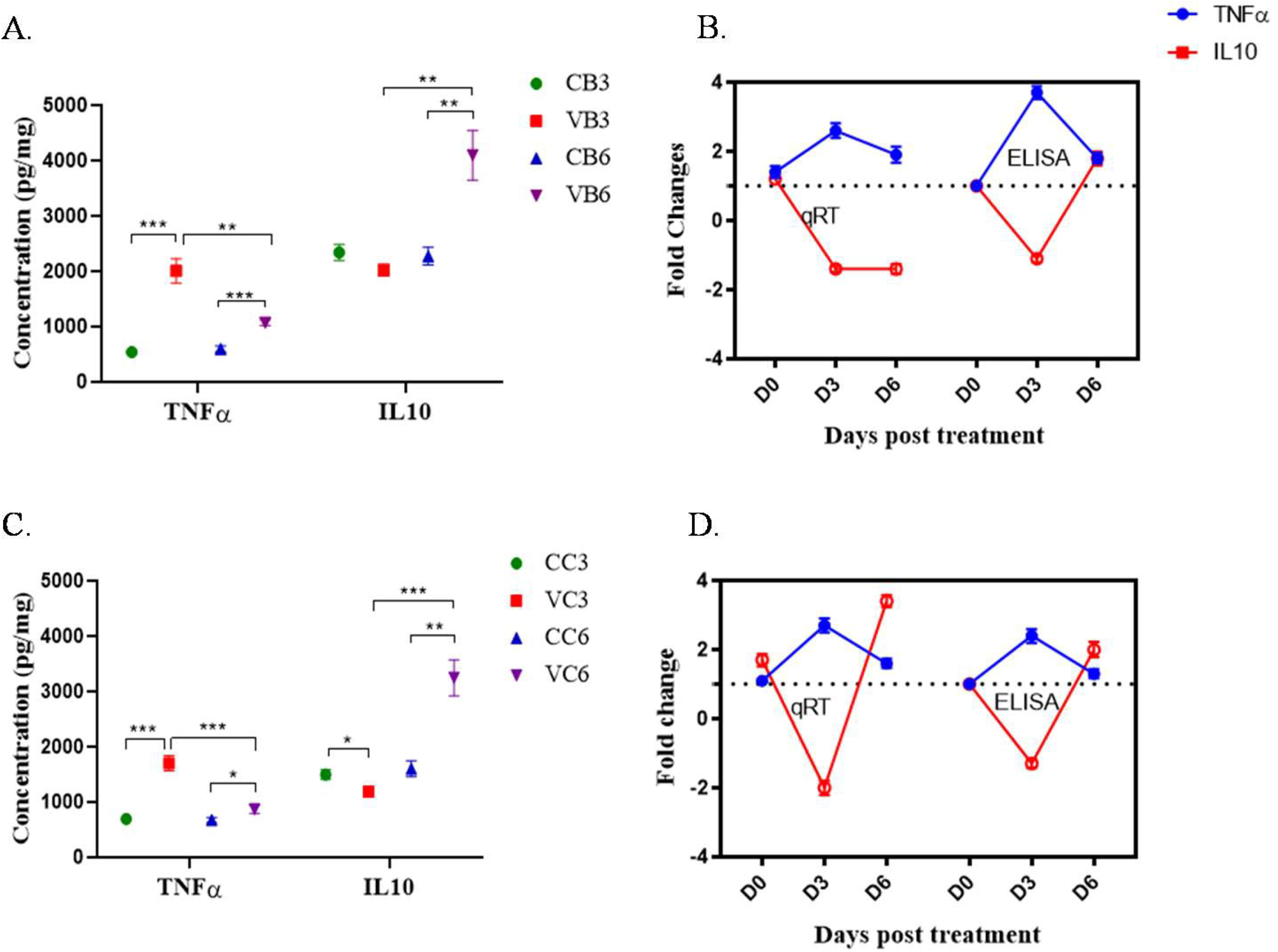
Protein level gene expression and comparative analysis of qRT PCR and ELISA data for TNFα and IL10. Mean values (n=3) of protein level concentration (in pg mg ^-1^) with standard deviation of TNFα (blue) and IL10 (red) expression on days 0, 3 and 6 for control and vancomycin treated A. BALB/c (CB and VB) and C. C57BL/6 (CC and VC) are shown. Statistical significance was calculated by two-way ANOVA (‘***’ corresponds to P ≤ 0.001, ‘**’ corresponds to P ≤ 0.01, ‘*’ corresponds to P ≤ 0.05 level of significance). Fold change values of expression of TNFα (blue) and IL10 (red) to compare the values obtained from qRT-PCR and ELISA studies are shown for B. BALB/c, and D. C57BL/6. Error bars of the data are already shown in preceding figures.

### Effect of vancomycin treatment on gut barrier integrity

Previous reports stated that gut microbiota is responsible to maintain the gut barrier integrity(Ulluwishewa *et al*., 2011; Feng *et al*., 2019). Alhough the effect of microbiota perturbation on gut barrier integrity of different immune biased mice (Th1 and Th2) is not clear. The current results revealed that treatment, with vancomycin for three days, massively disrupts the gut barrier integrity as was evident from FITC-dextran based gut permeability assay. Serum FITC-dextran level was significantly higher in day 3 treated mice compared to the day 0 control mice. But surprisingly the level decreased to normal (day 0 control) on the sixth day of treatment in both mice strains (Figure 5C). These results prompted us to evaluate the gene expression of different colonic tight junction proteins (occludin and claudin1) that maintain the barrier function of gut (Chelakkot, Ghim and Ryu, 2018). Results revealed that the expression of claudin1 gene decreased continuously from day one to day six following treatment with vancomycin in BALB/c and C57BL/6 mice (Figures 5A and 5B). However, the expression of occludin gene decreased continuously from day zero to day six in BALB/c mice and from day zero to day four in C57BL/6 mice. On day five and six, in C57BL/6 mice, we observed a slight increase in occludin gene expression compared to its day three following vancomycin treatment.

**Figure 5.**
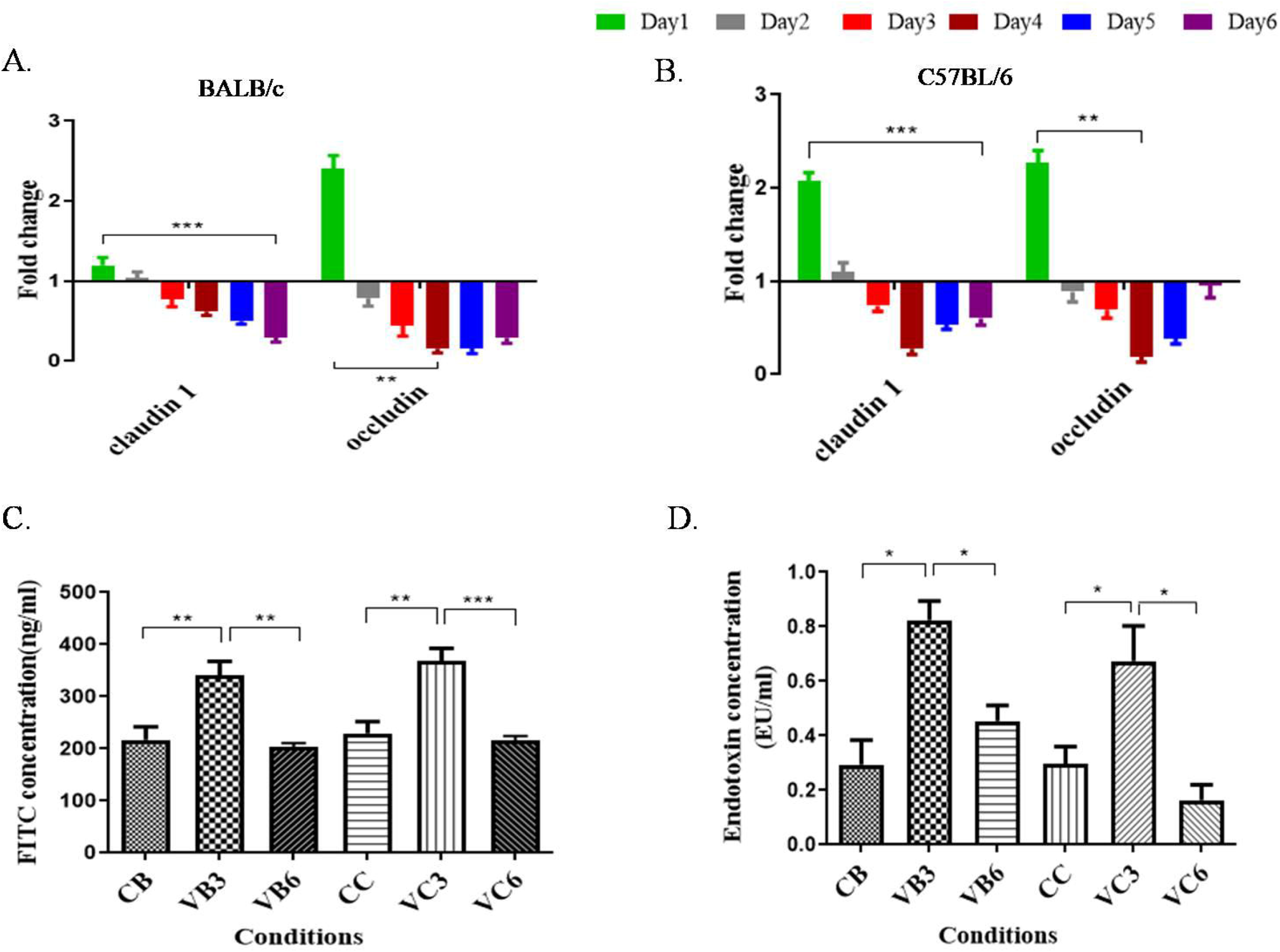
Measurement of intestinal integrity of BALB/c and C57BL/6 mice following treatment with Vancomycin. Transcriptional expression levels of tight junction genes, claudin 1 and occludin, in gut tissue by qRT-PCR are shown in vancomycin treated and untreated (control) groups for A. BALB/c and B. C57BL/6 mice C. Gut permeability data by measuring FITC dextran concentration in serum. D. Endotoxin concentration in the serum for both strains of mice are shown. where CB, VB3 and VB6 implies untreated (control), day 3 (D3) and day 6 (D6) post vancomycin treated BALB/c and CC, VC3 and VC6 denote the same for C57Bl/6 mice. The colors corresponding to different days are shown above the panels A. and B. Comparisons among the groups were calculated by two-way ANOVA. In the figure for panels ‘A.’, ‘B.’, ‘C.’. and ‘D.’, ‘***’ corresponds to P ≤ 0.001, ‘**’ corresponds to P ≤ 0.01, ‘*’ corresponds to P ≤ 0.05 level of significance.

Till day three following vancomycin treatment, both FITC dextran data and expression of tight junction genes showed almost similar results, i.e. decrease in gut barrier integrity. While on day six following vancomycin treatment, FITC-dextran studies suggested restoration of the gut barrier for both BALB/c and C57BL/6 mice, but the expression of claudin1 and occludin genes comparatively remained repressed with respect to their time-matched control mice. Further studies are required to understand this apparent discrepancy which may be due to a different mechanism, independent of these two genes that may exist to re-establish the gut barrier function.

In agreement with increased gut permeability, we intended to see whether it also induce the translocation of microbial products into the systemic circulation, as a result of barrier disruption. In both strains, serum endotoxin level was highest on the third day following vancomycin treatment compared to control mice and it was reduced on the sixth day of treatment (Figure 5D). On day six, BALB/c mice had slightly higher endotoxin level in serum compared to C57BL/6 mice. From the above findings, it is clear that the disruption of gut barrier integrity is strongly associated with the Proteobacteria level, whereas restoration is associated with Verrucomicrobia abundance in both mice strains

### Differential level of Verrucomicrobia in gut regulates blood glucose level following vancomycin treatment

Antibiotic mediated gut microbiota perturbation can affect different host metabolic functions, one such measurement involves the regulation of blood glucose homeostasis (Vrieze *et al*., 2014; Zarrinpar *et al*., 2018; Khan *et al*., 2019). Our results showed a high abundance of Verrucomicrobia phylum on the 6th day of vancomycin antibiotic treatment. Since previous studies reported that *Akkermansia* sp. from Verrucomicrobia phylum positively regulated glucose metabolism(Dao *et al*., 2016; Plovier *et al*., 2017) we intended to see how vancomycin induced time-dependent change in microbiota profile regulates blood glucose level. We performed an oral glucose tolerance test (OGTT) from 0 to 90 minutes in both strains of mice. From the glucose tolerance test, it was revealed that glucose metabolism was different in control and vancomycin treated mice (Figures 6A and 6B). Important to note that the results from OGTT studies for control animals for both BALB/c and C57BL/6 remain unchanged on days zero, three and six (data not shown). On the day third following vancomycin treatment, fasting blood glucose (0^th^ min) levels in the Th2- and Th1-biased mice (BALB/c 194.6± 6.3 mg dl ^-1^ and C57BL/6 186±6 mg dl ^-1^) were significantly higher than their respective zero-day untreated (BALB/c 115±3 mg dl ^-1^ and C57BL/6 126±4 mg dl ^-1^) mice. On day sixth following vancomycin treatment glucose levels dropped (BALB/c 148.6±7 mg dl ^-1^ and C57BL/6 103±5 mg dl ^-1^). The reduction of blood glucose level on day six following vancomycin treatment was more prominent in C57BL/6 compared to BALB/c mice (Figures 6A and 6B). The metabolism rate of glucose in the blood of the sixth-day vancomycin treated mice was faster than the third day treated mice. This rate was higher in vancomycin treated C57BL/6 mice compared to BALB/c mice. Next, we hypothesized that the differential level of Verrucomicrobia of C57BL/6 and BALB/c mice has an effect on the blood glucose level. To prove the causal role of Verrucomicrobia, we transplanted the cecal microbiota from sixth-day vancomycin treated mice (high Verrucomicrobia) to third-day vancomycin treated mice. We observed a significant drop in blood glucose level in the third-day vancomycin treated C57BL/6 mice after CMT. However, the blood glucose level remained unchanged even after CMT in the third-day vancomycin treated BALB/c mice (Figures 6A and 6B). These data together suggest that high Verrucomicrobia level on the sixth day treated C57BL/6 mice actually improves blood glucose level, which is not possible with a comparatively low level of Verrucomicrobia present in BALB/c mice following sixth-day vancomycin treatment.

**Figure 6.**
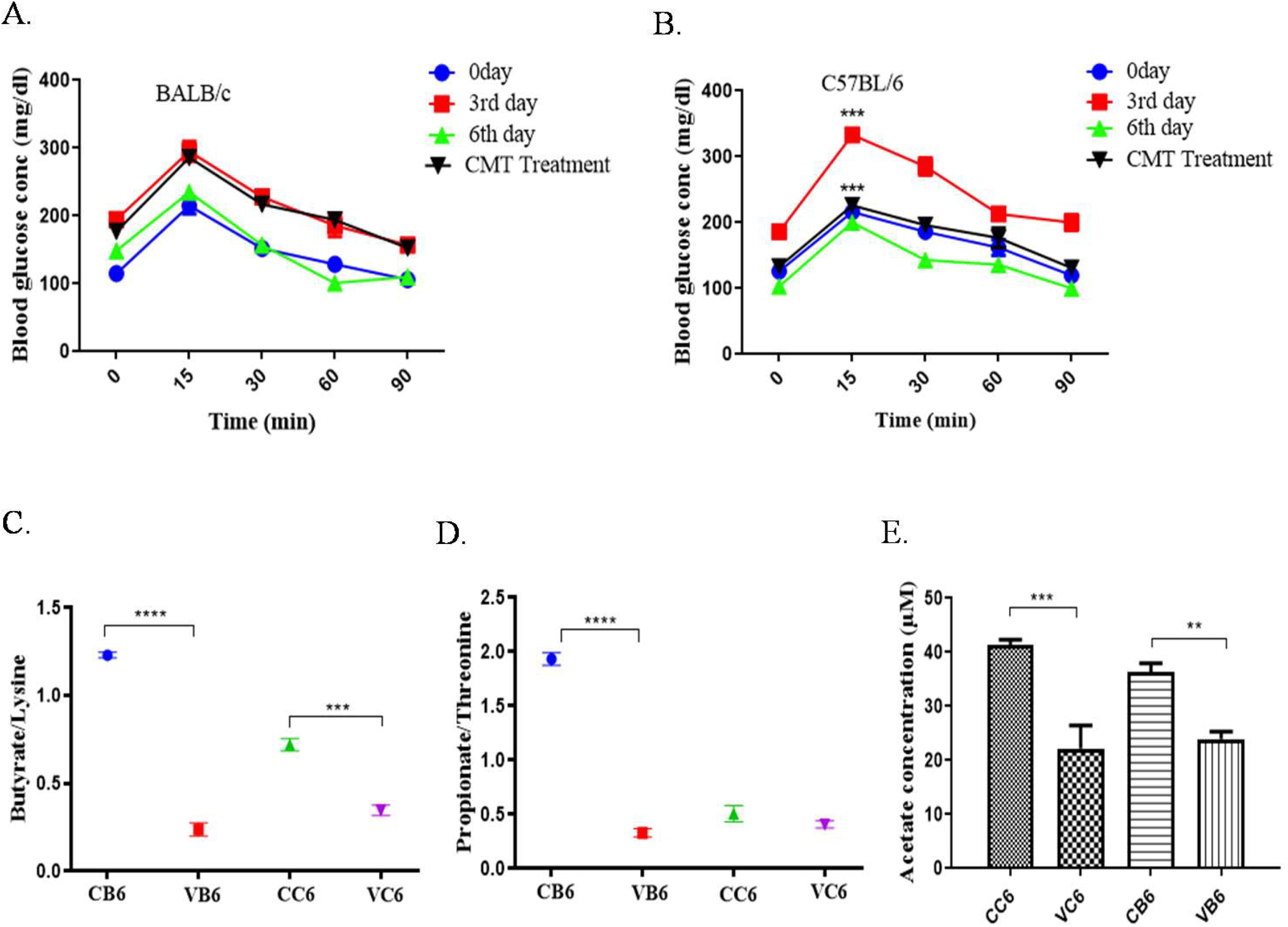
Glucose tolerance and abundance of select metabolites in serum. Kinetics of fasting blood sugar in A. BALB/c B. C57BL/6 mice following treatment with Vancomycin on days 0, 3 and 6 and following treatment with CMT from day 6 vancomycin treated mice transferred to vancomycin treated day 3 group of mice. Ratio of abundance, from chemometric^1^H-NMR studies for major short chain fatty acids, of C. butyrate production over lysine and D. propionate production over threonine in untreated control BALB/c (CB) and C57BL/6 (CC) and Vancomycin treated BALB/c (VB) and C57BL/6 (VC) are compared for Day 0 and Day 6 following treatment with vancomycin. In addition, E. acetate concentration in the serum by using acetate detection kit on day 6 in vancomycin treated groups of mice (VB6, VC6) along with the time matched control mice (CB6, CC6) of BALB/c and C57BL/6, respectively. In the figure, ‘****’ corresponds to P ≤ 0.0001, ‘***’ corresponds to P ≤ 0.001, ‘**’ corresponds to P ≤ 0.01, level of significance).

### Metabolites level in serum changes following vancomycin treatment

Antibiotic treatment can drastically reduce the Short-chain fatty acids (SCFAs) level, which are important regulators of several host physiological functions (Zarrinpar *et al*., 2018; Venegas *et al*., 2019). We measured the amount of SCFAs in the host serum using NMR based metabolomics. The current results showed that the ratios of the abundance of butyrate/lysine (Figure 6C) and propionate /threonine (Figure 6D) (Table.3) in serum were reduced significantly on day six following treatment with vancomycin. This might indicate the significant decrease in the conversion of SCFAs such as butyrate from lysine and propionate from threonine of vancomycin treated mice. The results further suggested the accumulation of the substrate (lysine, threonine) in blood and less detection of SCFA in the vancomycin treated day six group of mice (VB and VC on D6). We further measured the abundance of acetate in the serum of both BALB/c and C57BL/6 mice by using an acetate detection kit (Figure 6E). In both BALB/c and C57BL/6 mice, the concentrations of acetate on the day six following vancomycin treatment was found to be significantly lower than their respective time match untreated group of mice. Hence, these results indicate that the serum SCFA level decreased with vancomycin treatment and high Verrucomicrobia abundance on the 6th day cannot restore the SCFA level in both Th1 and Th2 biased mice.

### Effect of vancomycin treatment on metabolic hormones

Our previous results indicated that the rate of glucose metabolism and SCFA concentration in serum altered during vancomycin treatment. Circulating SCFAs are also related with different metabolic hormones(Larraufie *et al*., 2018; Müller *et al*., 2019) Considering that we further measured associated metabolic hormones such as insulin, Peptide tyrosine tyrosine (PYY) and leptin (regulates appetite) in mice during vancomycin treatment. Results revealed that insulin level decreased significantly on the day six compared to day three following treatment in C57BL/6 mice, but not in the BALB/c mice (Figure 7A). Hence, vancomycin treatment on day six showed a reduced amount of serum insulin concomitant with the blood glucose level in C57BL/6 mice. However, insulin levels on day three following vancomycin treatment were significantly higher compared to their respective day zero untreated group of mice for both strains. No significant changes were observed in serum leptin concentration during vancomycin treatment in both BALB/c and C57BL/6 mice with respect to their respective untreated controls (Figure 7B). Further results revealed that the concentration of PYY hormone in the gut decreased in both BALB/c and C57BL/6 mice from day zero to day six following vancomycin treatment (Figure 7C) Hence, our results suggest that vancomycin mediated gut microbiota perturbation may regulate blood glucose and insulin level differentially between C57BL/6 and BALB/c mice in a time-dependent manner.

**Figure 7.**
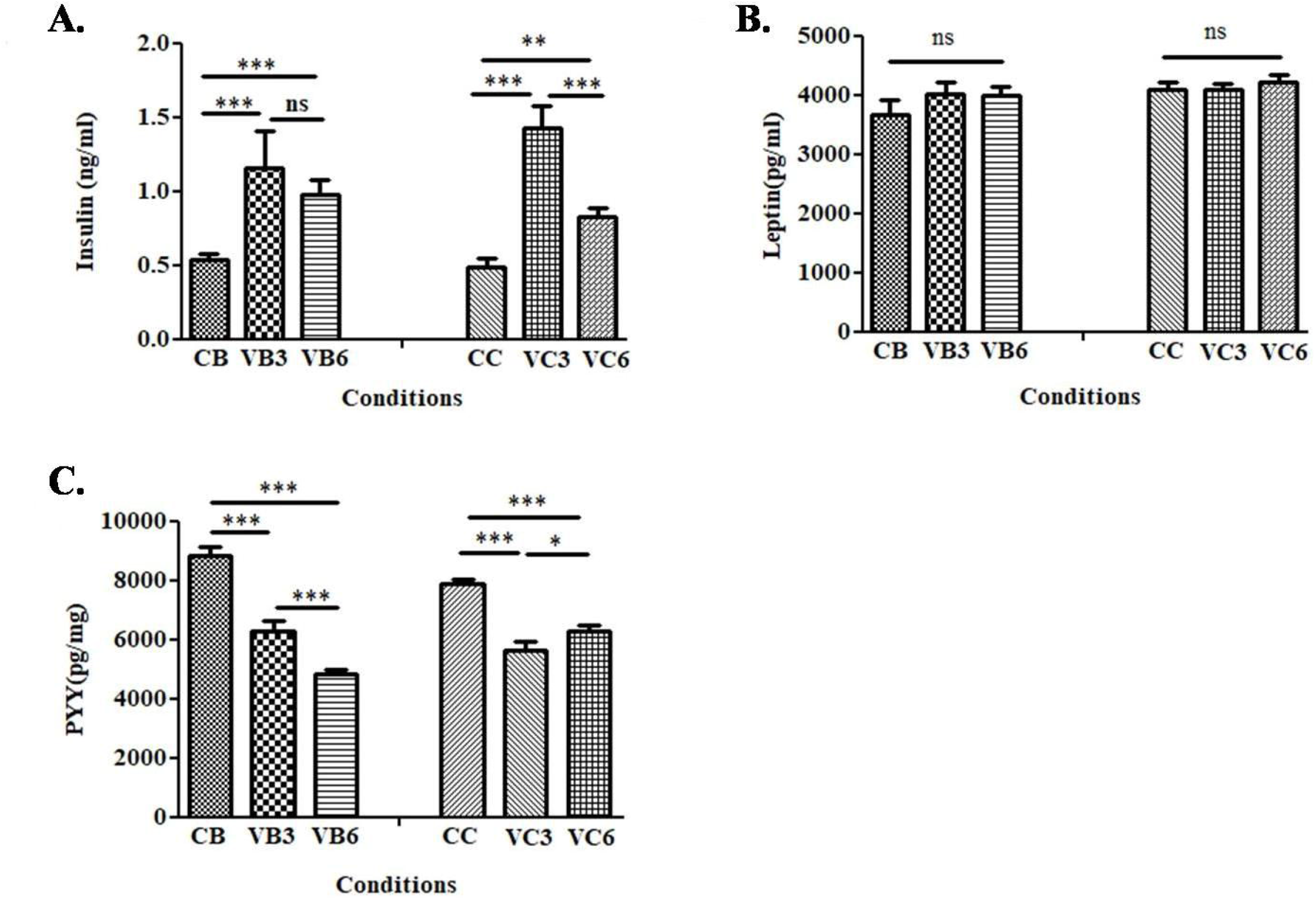
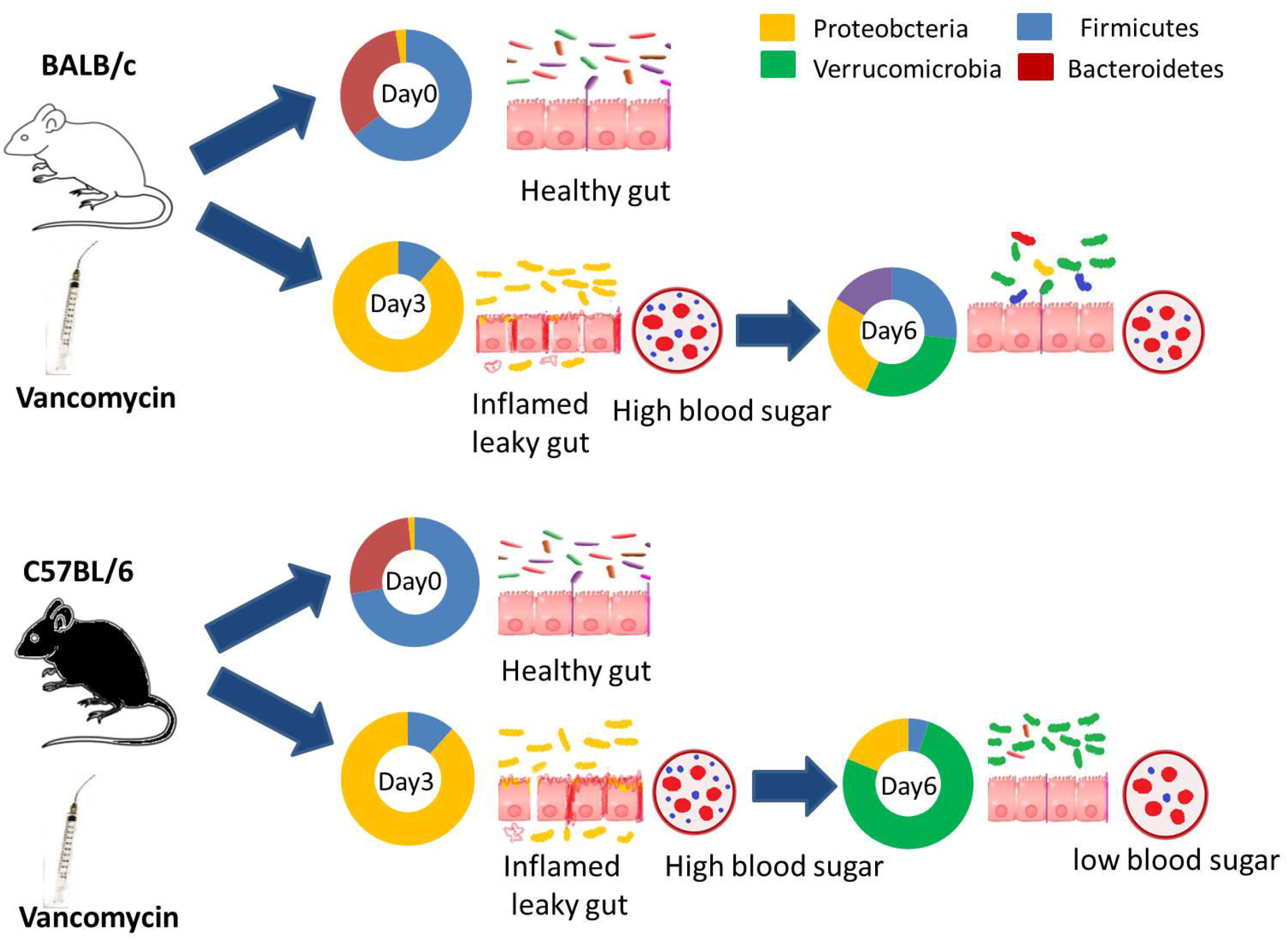
Changes in select hormones in serum. Abundance of A. Insulin (ng ml^-1^), B. Leptin (ng ml^-1^) and C. PYY (pg mg ^-1^) in the serum of control BALB/c (CB) or C57BL/6 (CC) and vancomycin treated mice on third day (VB3, VC3) and sixth day (VB6, VC6) of BALB/c and C57BL/6 mice respectively. Comparisons among the groups were calculated with two-way ANOVA. In the figure, ‘***’ corresponds to P ≤ 0.001, ‘**’ corresponds to P ≤ 0.01, ‘*’ corresponds to P ≤ 0.05 level of significance). Error bars shown are one standard deviation from the mean value of four replicates (n=4).

## Discussion

It was reported earlier that the treatment with vancomycin reduced the abundance and diversity of the gut microbiota (Vrieze *et al*., 2014; Isaac *et al*., 2016; Sun *et al*., 2019). The current study corroborated with the fact that mice treated with vancomycin had decreased levels of Firmicutes and Bacteroidetes and elevated levels of Proteobacteria in the initial four days. It is apparent that the increase in the one phylum would repress the abundance of other phyla and as a sequel, the overall diversity of the gut microbiota would decrease (Mosca, Leclerc and Hugot, 2016). In the current study, we observed an increase in only Proteobacteria phylum caused a decrease in the diversity of gut microbiota. Escherichia and Shigella genera of Proteobacteria phylum belong to the Gram-negative group of bacteria which caused an increase in endotoxin level in the blood through their LPS (Steimle, Autenrieth and Frick, 2016). These bacteria activate the TLR4 receptor present in the gut epithelial cell, and this causes increases in the expression of pro-inflammatory cytokines (Akira and Hemmi, 2003; Rallabhandi *et al*., 2008). In the current study, concerted effects of the increase in pathogenic Proteobacteria and a decrease in Firmicutes in the gut caused increased inflammation and endotoxin level of mice during initial days of vancomycin treatment. Firmicutes, specifically the Clostridium group present in the gut, produces short-chain fatty acids like acetate, butyrate, propionate from complex carbohydrate foods (Venegas *et al*., 2019). Bacteria belong to Instentimonas (Firmicutes phylum), produce butyrate from lysine and Bacteroidetes produces propionate from threonine in the gut (Bui *et al*., 2015; Neis, Dejong and Rensen, 2015). The production of these SCFAs in the gut suppresses the LPS and pro-inflammatory cytokines like TNF-α, IL-6 level and enhances the release of the anti-inflammatory cytokine like IL10 (Vinolo *et al*., 2011; Morrison and Preston, 2016). The current study revealed a decrease in Firmicutes and Bacteroidetes levels following treatment with Vancomycin till day four. The decrease, in the levels of Firmicutes and Bacteroidetes, may result in less yield of SCFAs and higher production of inflammatory cytokines in mice.

It is well established that over-expression of Inflammatory cytokine like TNFα is associated with the higher gut permeability by suppressing the expression of tight junction proteins like occludin and claudin 1 (Al-Sadi *et al*., 2013; Rios-Arce *et al*., 2017). Initial days of vancomycin treatment caused the lower expression of tight junction protein resulting in increased gut permeability. During obesity and diabetic condition, metabolic endotoxemia had been observed where endotoxin (LPS) level increased in the blood to cause inflammation and impaired glucose metabolism of the host (Hawkesworth *et al*., 2013; Boutagy *et al*., 2016). The current report revealed a higher concentration of fasting insulin and glucose in the serum of BALB/c and C57BL/6 mice on day 3 following vancomycin treatment compared to the control mice. The increased glucose and serum insulin level hinted at insulin resistance. The changes, in glucose and insulin levels, were associated with the higher abundance of Proteobacteria and endotoxin level on the day three following vancomycin treatment in both strains of mice.

The changes, in gut microbiome profile till day 4, were different from post day four of treatment, with Vancomycin. The profile of the changes in gut microbiota was distinct and varied between C57Bl/6 and BALB/c mice. It was reported that an increased abundance of Verrucomicrobia caused a decrease in inflammation and enhanced glucose metabolism of the host(Fujio-Vejar *et al*., 2017; Plovier *et al*., 2017; FUJISAKA *et al*., 2018). On the sixth day of vancomycin treatment, in the current study, a decrease in Proteobacteria and increase in Verrucomicrobia caused an anti-inflammatory effect and enhanced glucose metabolism in the gut. Specifically in C57BL/6 mice, with a significant decrease in Proteobacteria and increase in Verrucomicrobia phylum on day six following vancomycin treatment caused decreased tlr4 expression and increased tlr2 expression in the gut. However, on the sixth day following vancomycin treatment, replacement of Proteobacteria by Verrucomicrobia caused significant improvement in glucose metabolism of mice by bringing back the fasting glucose and insulin level in the blood to normal level. Following a successful transfer of the cecal sample from the sixth day of vancomycin treated C57BL/6 mice (*A. muciniphila* level is above 70%) to the third day of vancomycin treated mice caused a significant decrease in blood glucose level of third-day vancomycin treated mice. The glucose level decrease following cecal microbiota transplant was comparable to the control glucose level. Since sixth-day vancomycin treated mice had a significantly higher level of Akkermansia, the current report prospectively hinting at the effective causal role of *A. muciniphila* in controlling the blood glucose level. On the sixth day of vancomycin treatment, *A. muciniphila* level was significantly higher in C57BL/6 mice than BALB/c. This higher level of Akkermansia is a very good supportive evidence of our suggestion of the role of *A. muciniphila* in reversing the glucose level to normal. This higher level of Verrucomicrobia in C57BL/6 could have caused a more prominent effect in decreasing glucose level and increasing insulin sensitivity in the blood of C57BL/6 than BALB/c mice.

Reports also suggested that higher abundance or production of SCFA usually leads to anti-inflammatory response (Vinolo *et al*., 2011). The current study further confirmed the proposition by showing higher SCFA yield in C57BL/6 than BALB/c. This observation is also in corroboration with the Th1-bias of C57BL/6 over Th2-immune bias of BALB/c mice. Reports also revealed that SCFA could stimulate PYY hormone (explain briefly the function or importance of PYY) production by activating Gq-coupled receptor, FFAR2, of endocrine cells present in the gut (Cahill *et al*., 2014; Larraufie *et al*., 2018). The current results revealed that following treatment with vancomycin the level of SCFA decreased. The decrease in SCFA level is concomitant with the reduction in the production of serum PYY. The observations so far prompted us to conclude the following proposition.

Host genetics is one of the major factors that regulate the gut microbiota composition and ecosystem (Korach-Rechtman *et al*., 2019). C57BL/6 and BALB/c are two genetically different inbred mouse lines, which are respectively Th1 and Th2 immune biased mouse strains and differ in their baseline microbiota composition (Watanabe *et al*., 2004; Fransen *et al*., 2015). Here we found that the gut microbial population responds differentially against the vancomycin challenge, which is associated with higher a) abundance of Verrucomicrobia and b) production of SCFAs in C57BL/6 compared to BALB/c mice. The changes in the gut microbiota through vancomycin perturbation can alter host metabolism like glucose tolerance significantly between two strains of mice. Overall, the time-dependent perturbation of gut microbiota by vancomycin was not random. It followed a particular pattern. It affects the host in two different ways; Initial doses caused increased in pathogenic bacteria in the gut which caused a most deleterious effect on the host while continued later doses of vancomycin caused in increased Verrucomicrobia phylum in the gut which showed some beneficial effects on the host.

## Author Contribution Statement

PR performed all experiments and drafted the manuscript. PR and PA designed the experiments. UP designed critical parts of some experiments and also contributed to the manuscript preparation. PA conceptualized, supervised the studies and finalized the manuscript.

## Acknowledgement

This research received no specific grant from any funding agency in the public, commercial, or not-for-profit sectors. Authors extend their thanks to the NMR facility of NISER for assisting in collecting 1D proton-NMR FIDs and also thank the animal facility of NISER for their assistance. This manuscript has been released as a pre-print at BioRxiv (https://www.biorxiv.org/content/10.1101/516898v2).

## Conflict of Interest

The authors declare that there is no conflict of interest.

## Funding and payment

The current work (necessary resources to perform the experiment and the infra-structure for the laboratory) was supported by the parent institute National Institute of Science Education and Research through intramural funding by DAE, GoI India. The current work was not supported through any extra-mural funding except the PhD fellowship to PR by the Council of Scientific and Industrial Research (CSIR), Govt. of India, India. The current authors have no support to pay for the open-access or article processing fees to publish this research article.

